# Open source software tool for the automated detection and characterization of epithelial cells from trace biological samples

**DOI:** 10.1101/2020.03.24.006296

**Authors:** Anita Olsen, Mekhi Miller, Vamsi K. Yadavalli, Christopher J. Ehrhardt

## Abstract

This paper presents a strategy for an unsupervised workflow for identifying epithelial cells in microscopic images and characterizing their morphological and/or optical properties. The proposed method can be used on cells that have been stained with fluorescent dyes and imaged using conventional optical microscopes. The workflow was tested on cell populations that were imaged directly on touch/contact surfaces and stained with nucleic acid dyes to visualize genetic content. Our results show that this approach could be a useful strategy for characterizing differences in staining efficiency and/or morphological properties of individual cells or aggregate populations within a biological sample. Further, they can potentially reduce the laborious nature of microscopic analysis and increase throughput and reproducibility of similar studies.

## Introduction

Characterizing the cellular components of trace biological evidence and its relationship to genetic material within the sample has the potential to address long standing questions surrounding the deposition, persistence, and transfer of DNA in evidentiary material. Such analysis is particularly relevant for samples generated from touch/contact. Toward this end, recent studies have utilized the microscopic imaging (typically optical microscopy) of cells combined with fluorescence staining of nucleic acids to examine correlations between the optical properties of cells and DNA yield (1), analyze contributor-specific patterns in DNA yield (2), or to detect biological material in various types of touch/contact evidence samples (3,4). In nearly all of these examples, the detection and enumeration of the epithelial cells in the imaging data was performed manually (i.e., through visual inspection/identification). This can be time-consuming, laborious and reduce throughput. Further, reproducibility of results across laboratories or even analysts is challenging.

To address this, we have developed an analysis pipeline for the automated processing of microscopic images using the open-source software ‘Cell Profiler’ (5,6). This pipeline was specifically optimized for epithelial cell populations deposited in touch or trace biological samples. Although there are several software tools for the unsupervised recognition and segmentation of cells from microscopic images, most rely on nuclear staining and/or counterstaining of different cellular constituents. For touch/contact cell populations, a vast majority of cells are likely to be anucleate and to have limited amounts of both intracellular DNA and other traditional targets for fluorescent probes (7,8). This may limit the effectiveness of existing detection methods.

In order to make this process universal, the input for our analysis pipeline can be microscope images in any standard digital format (e.g. .jpg, .tiff, etc.). These images contain cells that have been stained with a fluorescent DNA dye. The script detects and quantifies cells within the image and then extracts a series of morphological and optical measurements from each cell. Since recent work has shown that fluorescence and/or structural properties of cells can indicate tissue origin and contributors (9,10), this approach may be used to indicate a variety of forensically relevant attributes from the sample. We demonstrate the potential applications of this technique by analyzing microscopic images of trace epithelial cell samples generated from 15 different donors and characterizing differences in cell attributes across contributor cell populations.

## Methods

The sample set consists of epithelial cell samples collected from 15 individuals following Institutional Review Board (IRB) approved protocol (ID# HM20000454_CR5). Touch epidermal cells were obtained by having individuals press their finger or thumb directly onto a glass slide surface for 10 seconds. For nucleic acid staining, GelGreen® Dye was used. The choice of this dye was based on sensitivity (11), as well as other attributes such as low toxicity and improved stability at room temperature. Cell populations deposited onto each slide were stained with GelGreen (P/N 41005; Biotium, Inc.) following manufacturer’s protocols. Briefly, 200 μL of a 3X dilution in sterile H_2_O was applied to the slide and then incubated at 40°C for 30 minutes. Following this, slides were rinsed with 2 mL of sterile H_2_O and stored at dry at room temperature until analysis.

Fluorescence images were taken across print samples for 10-30 cells per contributor cell population. Since touch/contact epidermal cells can present autofluorescence (1,9), images of untreated cells were also collected as a control. The analysis pipeline was created using the CellProfiler software for detecting individual cells and defining their boundaries based on fluorescence signal (5,6). Cells were identified and masked via fluorescence intensities above background, and contamination was filtered out by size. Once cells were identified and their area defined spatially within the image, other morphological and/or fluorescence measurements were collected (e.g., circularity, aspect ratio, diameter). The entire software pipeline and list of measurement variables collected is available as supplementary material (12).

## Results and Discussion

Cell populations from fingerprints representing 15 different contributors were imaged using both brightfield and fluorescence microscopy. Detection and segmentation of cells was based on the fluorescence image (**Figure 1**). The analysis pipeline for detecting and segmenting cells from each image was optimized to such that smaller debris/detritus can be omitted based on size. The total number of cells detected and analyzed from each contributor fingerprint ranged between 31 and 174 (**Table 1**). Generally, cells ranged between ~20μm and ~80μm in diameter. This is consistent with dimensions of epithelial cells derived from the outermost layer of the epidermis (i.e., corneocytes). After segmenting cells from the image, measurements were automatically generated from individual cells using 33 different variables representing different aspects of both fluorescence intensity across the cell surface and size/morphology. Data for four of those measurements (cell area, median fluorescence, form factor, and MAD intensity) across contributor cell populations are shown in **Figure 2**. Interestingly, although cell area showed largely overlapping distributions across contributor cell populations (**Figure 2a**), the median fluorescence intensity was markedly lower in Individuals 3 and 14 compared to other contributors. Similarly, values for the form factor variable, a measurement which captures the circularity of a cell (4π*Area/Perimeter^2^), was significantly lower for cell populations from Individuals 3 and 14 compared to other contributors (**Figure 2c**). A comparison of median absolute deviation intensity, a measure of the spatial heterogeneity of fluorescence across the cell (13), showed overlapping ranges of values for most contributor cell populations with the exception of Individual 14 who exhibited a lower median value (**Figure 2d**).

**Table 1.** Cells imaged for contributor populations

| Individual | 1 | 2 | 3 | 4 | 5 | 6 | 7 | 8 | 9 | 10 | 11 | 12 | 13 | 14 | 15 |
| --- | --- | --- | --- | --- | --- | --- | --- | --- | --- | --- | --- | --- | --- | --- | --- |
| # Cells imaged | 82 | 72 | 131 | 59 | 60 | 69 | 93 | 174 | 31 | 38 | 66 | 162 | 44 | 121 | 145 |

**Figure 1.**
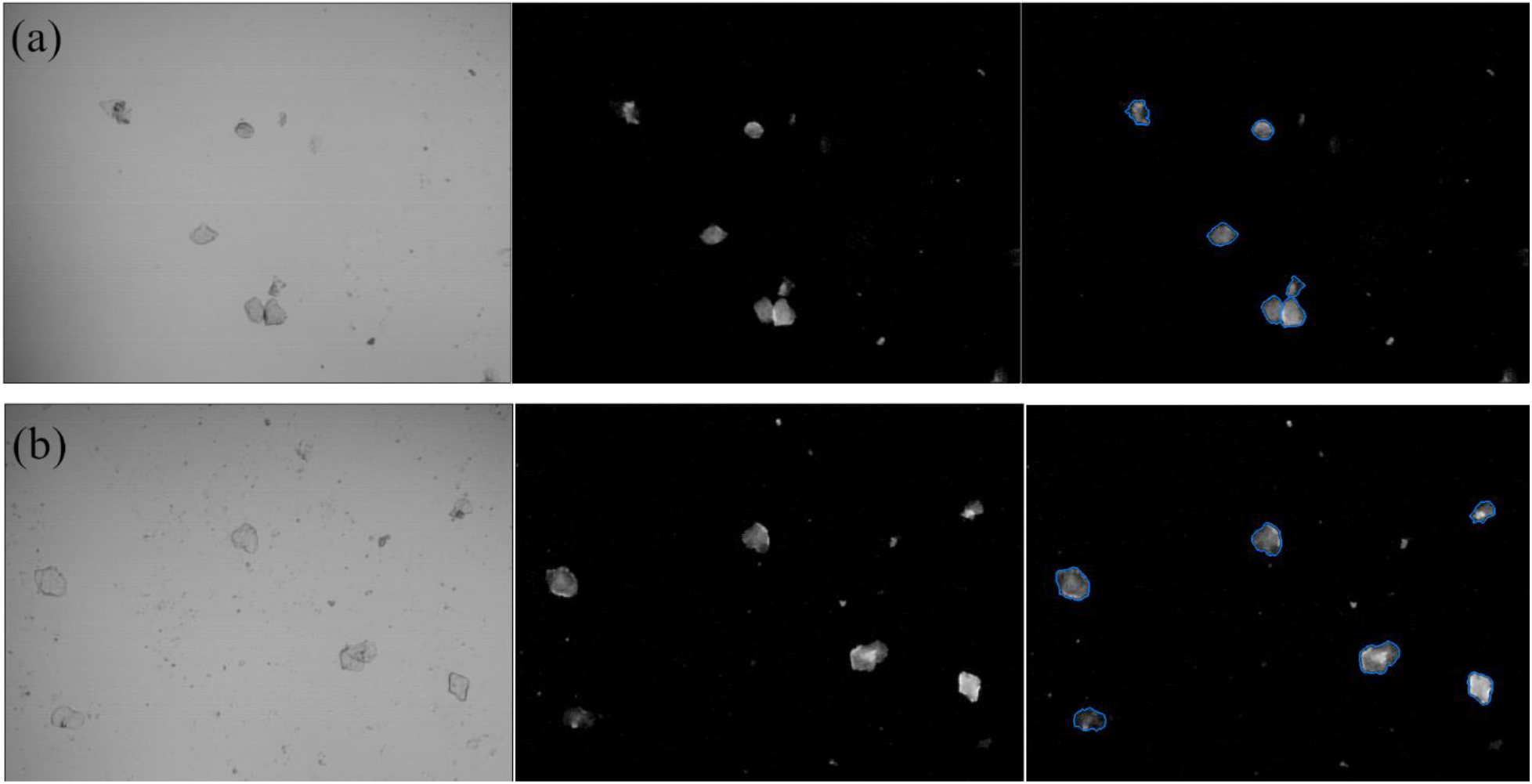
Image processing workflow for cell populations from Individuals 6 (a) and 15 (b). Brightfield and fluorescence images are shown for the same field of view (left and middle panels respectively). Segmentation scripts are then applied to fluorescence images to detect individual cells and extract measurements (right panel).

**Figure 2.**
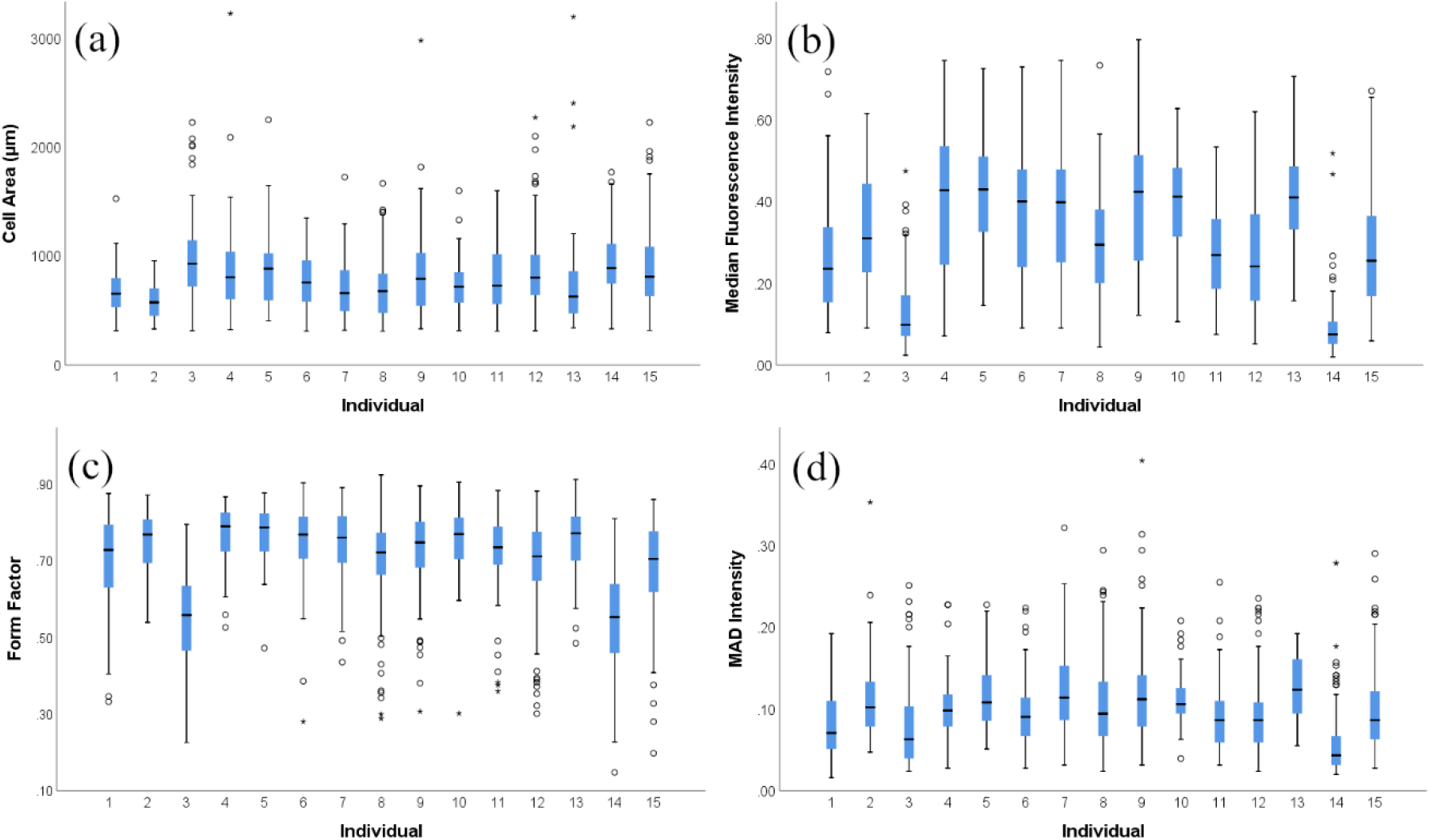
Comparisons of cell populations from different contributors for cell area (a), median fluorescence (b), form factor (c), and median absolute deviation (MAD) Intensity (d). Measurements are made on individual cells within each population.

As a complementary approach, multivariate analysis was performed using measurement data across all variables. The two-dimensional discriminant function plot shows that cell populations from Individuals 3 and 14 were easily differentiated from all other contributor cell populations (**Figure 3**). Additionally, a subpopulation of cells from Individual 1 were separated from all other contributor cell populations (top portion of plot in **Figure 3**). This observation that trace cell populations from certain individuals may have distinct morphological and/or structural properties is consistent with previous studies (9,14). Additionally, although contributor-specific trends in the deposition of cellular material and DNA have been observed (2,4), the propensity of individuals to systematically deposit more or less DNA in a touch/contact sample (i.e., shedder status) remains the subject of significant debate within the forensic community (7). This workflow which can quantify fluorescence on a *per cell* basis after treatment with a DNA-specific dye can therefore potentially be used to experimentally test hypotheses around shedder status and elucidate biological mechanisms governing the deposition, transfer, and persistence of trace DNA in forensic samples.

**Figure 3.**
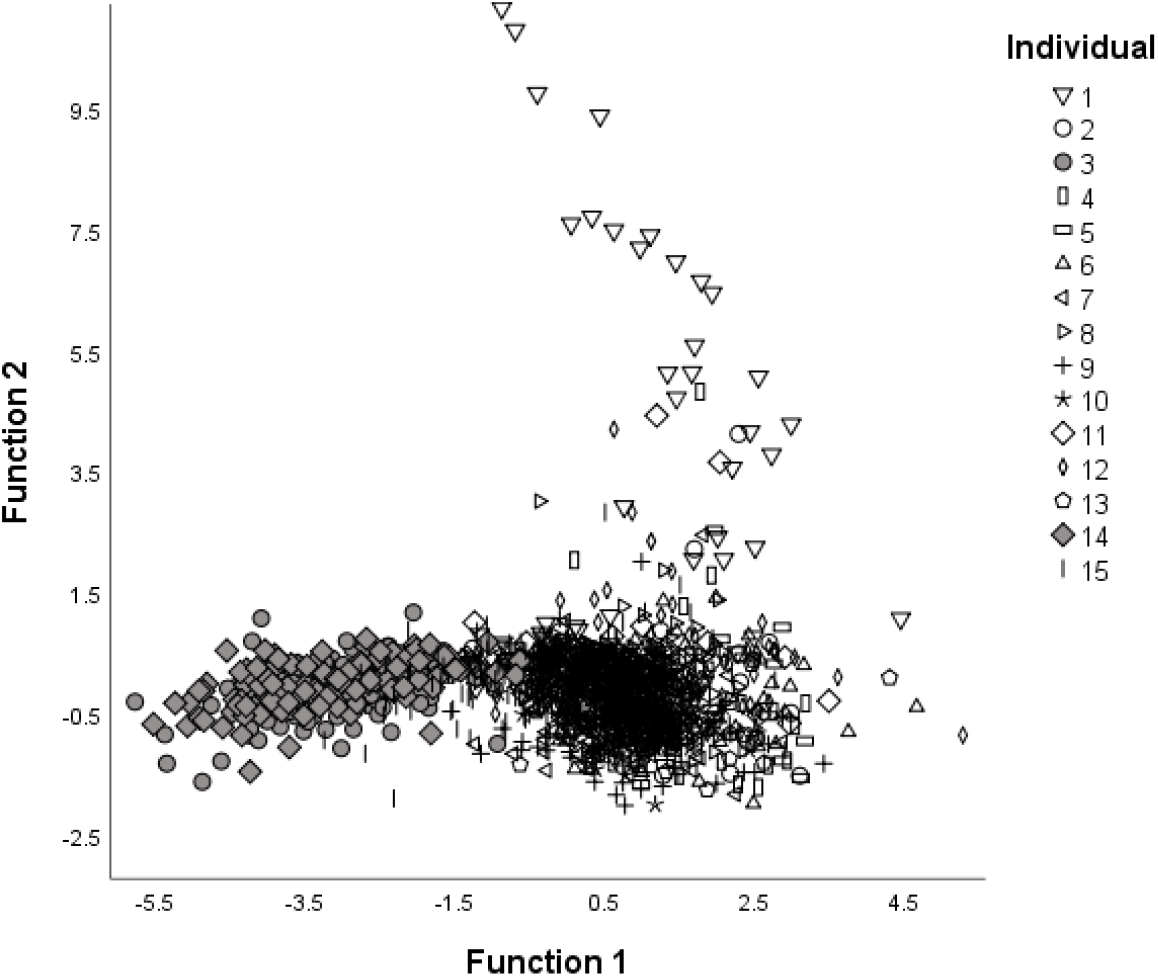
Multivariate analysis of morphological and fluorescence measurements across contributor cell populations. Cell populations 3 and 14 are clustered on the left side of the plot are shown with filled gray symbols (circle and diamond, respectively).

## Conclusions

In summary, we present an unsupervised workflow for the potentially high-throughput characterization of cell samples. Open source software is used (viz. CellProfiler) making this an easily accessible, low-cost tool. In our sample collection, ~1350 cells were collected across 15 different donors and subpopulations of individual donors could be identified. By staining the cells with fluorescence dyes, it is possible to quantify fluorescence on a *per cell* basis. This technique can therefore be used to investigate individual shedder status while also observing fundamental biological mechanisms governing the deposition, transfer, and persistence of trace DNA in forensic samples.

## Grant information

This project was funded by the National Institute of Justice Award numbers 2017-DN-BX-0186 (PI: Yadavalli).

## Competing interests

No competing interests were disclosed.

## Notes

https://doi.org/10.6084/m9.figshare.9947321.v1

